# Increased region of surround stimulation enhances contextual feedback and feedforward processing in human V1

**DOI:** 10.1101/2021.02.27.433018

**Authors:** Yulia Revina, Lucy S. Petro, Cristina B. Denk-Florea, Isa S. Rao, Lars Muckli

## Abstract

The majority of synaptic inputs to the primary visual cortex (V1) are non-feedforward, instead originating from local and anatomical feedback connections. Animal electrophysiology experiments show that feedback signals originating from higher visual areas with larger receptive fields modulate the surround receptive fields of V1 neurons. Theories of cortical processing propose various roles for feedback and feedforward processing, but systematically investigating their independent contributions to cortical processing is challenging because feedback and feedforward processes coexist even in single neurons. Capitalising on the larger receptive fields of higher visual areas compared to primary visual cortex (V1), we used an occlusion paradigm that isolates top-down influences from feedforward processing. We utilised functional magnetic resonance imaging (fMRI) and multi-voxel pattern analysis methods in humans viewing natural scene images. We parametrically measured how the availability of contextual information determines the presence of detectable feedback information in non-stimulated V1, and how feedback information interacts with feedforward processing. We show that increasing the visibility of the contextual surround increases scene-specific feedback information, and that this contextual feedback enhances feedforward information. Our findings are in line with theories that cortical feedback signals transmit internal models of predicted inputs.

**Significance Statement:** The visual system has circuit mechanisms for processing scene context. These circuits involve lateral and feedback inputs to neurons. These inputs interact with feedforward inputs and modulate neuronal responses to visual stimuli presented outside their receptive fields. Systematically investigating independent contributions of feedback and feedforward processes is challenging because they coexist even in single neurons. Here we use an occlusion paradigm to isolate feedback and lateral signals in human participants viewing natural scene images in fMRI. We show that increasing the visibility of the contextual surround increases scene-specific feedback information, which also enhances feedforward signals. Our findings are in line with theories that cortical feedback signals carry abstract internal models that combine with more detailed representations in primary visual cortex.

## Introduction

Sensory stimulation triggers a cascade of processing in a hierarchy of visual areas. This feedforward processing meets recurrent activity from the previous sensory input and triggers recurrent activity that will meet the next expected visual input. Recurrent processing contextualises and predicts the incoming signal and updates internal models and future recurrent streams. The contextualisation of feedforward information by feedback signals is essential for our understanding of cortical processing (Gilbert and Li, 2011). We know from animal recordings that cortical neurons are contextually modulated when their response to a feedforward stimulus feature is modified by the presence of surrounding features (Sugita, 1999; Shushruth, 2011). In visual cortex, this contextual information can be located far in the surround of a neuron’s receptive field. Consequently, contextual modulation of neurons is exerted by cortical feedback and lateral inputs (Angelucci, 2002). Cortical feedback inputs, at least in non-human primate cortex, arrive to discrete portions of cortical pyramidal neurons; mainly to the apical dendrites that branch up to layer 1 (Douglas and Martin, 2007). Feedback inputs are therefore computationally distinct from feedforward inputs arriving to basal dendrites. Recent conceptual shifts in our understanding of neuronal computation are contributing to a developing perspective on the significance of cortical feedback inputs in determining neuronal information processing (Larkum, 2013). This perspective requires techniques to probe brain processing that detect neuronal inputs, advancing previous studies that mainly measure neuronal outputs (i.e. spiking activity Larkum et al., 2018; Muckli et al., 2015). Functional magnetic resonance imaging (fMRI) is one such technique that detects pre- and postsynaptic inputs, offering a means to measure contextual feedback information to a region of cortex.

Understanding the nature of contextual modulation transmitted by cortical feedback and lateral interaction is vital for understanding the brain in behavioural and cognitive contexts (Gilbert and Sigman, 2007). This importance of cortical feedback and lateral interaction arises because contextual modulations on a neuron include influences from higher-level top-down processes including expectation, prior experience and goal-directed behaviour, which originate in higher cortical areas (Muckli and Petro, 2013). Therefore, describing neuronal substrates of cognition in brain networks including sensory areas requires us to measure not only stimulus-driven neuronal responses under discrete states of top-down influences (e.g. attention, expectation, task, working memory), but also feedback-driven responses in isolation from feedforward processing. Measuring feedback-driven modulations separate from stimulus-driven activity allows us to investigate the information contained in top-down influences. These signals alter neuronal responses to stimuli (Li et al., 2004; Schwiedrzik and Freiwald, 2017; Petro and Muckli, 2018), which may depend on other state variables (e.g. being conscious, Philips et al., 2016), therefore functionally determining the brain’s response to its environment (Friston, 2010; Clark, 2015).

We used fMRI, a brain imaging measure of energy consumption, and multivoxel pattern analysis (MVPA) to investigate how global natural scene features contextually modulate human V1. Our approach complements non-classical receptive field studies in rodent and monkey cortex, that measure spikes in response to a feedforward stimulus relative to contextual surround stimulation. However, the proposed tuning to pre-and post-synaptic activity in apical dendrites that might be detectable by fMRI allows us to capitalise on a signal that might not always be available in sharp electrode electrophysiology, where the input at the apical dendrites might not lead to a change in spiking output. Using partially occluded images, we parametrically vary the amount of global contextual information that we provide and measure the resulting contextual feedback (and lateral interaction) information to V1 both in the absence of feedforward information, and when feedback is integrated with feedforward information. If global features in the surround contextually modulate human V1, we hypothesized that scene information in non-feedforward-stimulated V1 voxels should decrease with progressive masking of the surround, and increased surround stimulation should modulate detectable scene information even when V1 voxels receive feedforward stimulation.

## Materials and Methods

### Subjects

We compensated twenty-nine subjects from the University of Glasgow to participate in the experiment (n = 13 males; mean age: 24.28 years, range: 19-41 years). Subjects provided informed written consent and the experiment was approved by the local ethics committee at the University of Glasgow (CSE01063). We excluded subjects if their data was at chance level classification performance in at least one feedforward control condition (n = 5) or poorly aligned (anatomically) between functional runs (n = 3, see *Voxel Selection and Analysis*, indicating substantial body movement between scans). Below we report results from 21 subjects with stable classification in feedforward control conditions (n = 10 males; mean age: 25.29 years, range 19-41 years).

### Stimuli

#### Feedback vs Feedforward condition

We used occluded natural scene stimuli to investigate cortical feedback signals in the absence of feedforward stimulation (Smith and Muckli, 2010; Muckli et al., 2015; Revina et al., 2018; Morgan et al., 2019). For the feedback conditions, the lower right image quadrant was occluded by a white rectangle. Here we expect that the retinotopic region of V1 responding to the white portion of the image receives no meaningful feedforward input and only cortical feedback signals (and lateral inputs). The white rectangle was placed 0.5° of visual angle diagonally from the centre of the image and spanned 11.6° × 9.2°. In the so-called ‘feedforward’ conditions, the corresponding quadrant of the scene was shown; V1 voxels responding to the lower image quadrant in this condition contain a mixture of feedforward, lateral and feedback inputs.

#### Scenes

We used two natural scene images for each participant, as natural scenes induce a lot of contextual associations (Bar 2004). Each scene was 600 x 480 pixels and spanned 24° × 19.2° of visual angle. We did not normalize the images in terms of low-level visual features, such as luminance, contrast or energy at each spatial frequency because we wanted the scenes to look as natural as possible. Smith and Muckli (2010) previously showed that contextual feedback signals in V1 cannot be solely attributed to these low-level visual features.

To investigate the contribution of surrounding contextual information on the brain activity patterns corresponding to the lower right quadrant, we manipulated the visibility of the surrounding ¾ of the scene with a Gaussian aperture in each quadrant (“bubbles”, Gosselin and Schyns, 2001) of various sizes to reveal the scene and produce the following types of stimuli: ¼ (no surrounding scene shown), Small Bubbles (standard deviation [SD] = 50 × 32 pixels), Medium Bubbles (SD = 90 × 56 pixels), Large Bubbles (SD = 125 × 100 pixels) and Full (surround fully visible). The study consisted of four experiments, with each subject participating in only one (n = 6; n = 4; n = 6; n = 5 respectively). In each experiment, stimuli were shown in four (out of the five possible) different conditions (**Figure 1A**). In Experiment 1, we used stimuli in the Full Feedback occluded condition, ¼ feedforward, Small Bubbles feedforward and Medium Bubbles feedforward conditions. In Experiment 2, we replaced Small and Medium Bubbles with Large Bubbles and the Fully Visible scene. In Experiment 3, we added the Fully Visible scene to test whether more contextual feedback would be seen in the Small and Medium Bubbles conditions if participants were more familiar with the full scene. In Experiment 4, we tested the effect of reducing the surrounding information around the occluded region using Small, Medium and Large Bubbles feedback conditions.

**Figure 1.**
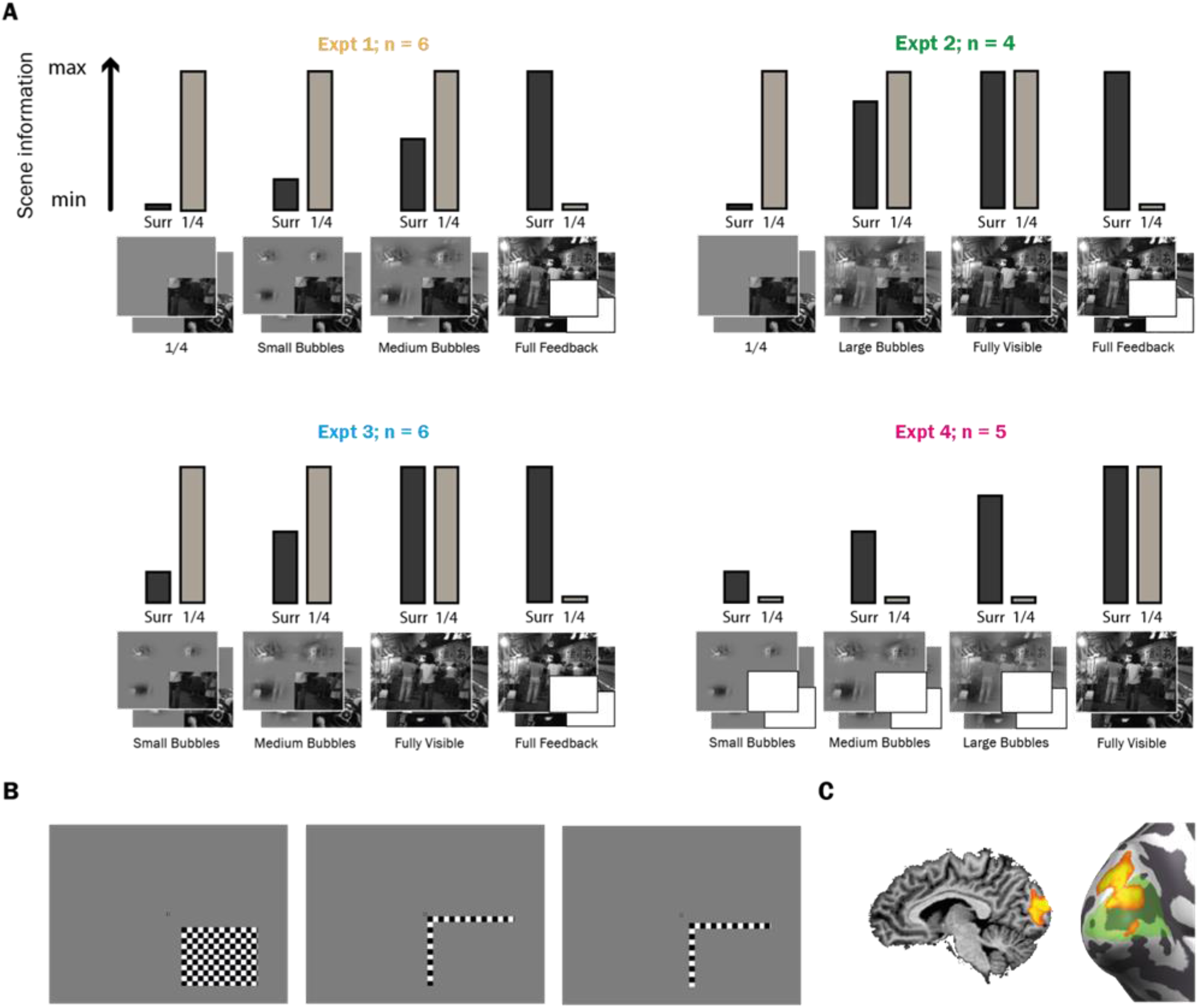
Stimuli. A) In feedback conditions the lower right image quadrant was occluded with a white rectangle, while in feedforward conditions the corresponding quadrant was visible. We manipulated scene visibility around the lower right quadrant with bubbles of various sizes to create 5 types of conditions: ¼, Small Bubbles, Medium Bubbles, Large Bubbles and Full. Dark bars labelled “Surr” illustrate the extent to which the surrounding ¾ of the scene was revealed. Light bars labelled “¼” illustrate the extent to which the lower right image quadrant was revealed. Bars are not to scale. B) Checkerboard stimuli were used to retinotopically map the occluded region in V1; left to right Target, Near Surround, Inner Border. C) The activation for the contrast of (Target – Near Surround) used to map non-stimulated V1 is shown on the occipital cortex on one subject, with V1 in green on the inflated visualization.

#### Occluded region mapping

We presented subjects with three contrast-reversing checkerboards (5 Hz) twice per run. The checkerboards either covered an inner rectangular part of the occluded region (*Target* – 2.5° diagonally from centre, 10.2° × 7.8° visual angle) or the border between the lower right quadrant and the rest of the stimulus (*Surround*). There were two types of surround checkerboard stimuli (**Figure 1B**) – *Near Surround* (0.5° diagonally from fixation, 11.6° × 9.2° visual angle) and *Inside Border* (1.5° diagonally from fixation). The activation in the early visual areas for the (*Target* – *Near Surround*) contrast is shown in **Figure 1C**.

### Experimental Design and Statistical Analysis

#### Task and procedure

We presented scenes on a uniform grey background using MRI compatible goggles (NordicNeuroLab) with 800 × 600 pixel screen resolution, which corresponded to 32° × 24° visual angle. In each experiment there were 8 types of trial (2 scenes in 4 different conditions). In each 12 second trial the stimulus was flashed on and off (200 ms on/ 200 ms off) 28 times (11.6 secs + variable fixation to account for uncertainty in timing). This flashing increases the signal to noise ratio compared to continuous presentation (Kay et al., 2008) and gives rise to a greater BOLD response (Boynton et al., 1996). Each trial type was presented sequentially, with the trial order randomized in each sequence. Each sequence lasted 96 seconds (8 × 12 s). A 12 second fixation period was included before and after each sequence of trials. Each experimental run lasted 10 min 48 seconds, containing four trial sequences and two mapping sequences (each mapping sequence consisted of *Target* and two *Surrounds*). There were four experimental runs in total. Thus, each stimulus was shown 16 times per experiment. Subjects’ task was to fixate on a central checkerboard and report a fixation colour change with a button press. Subjects pressed a different button depending on whether the colour change occurred during scene 1 or scene 2 (right index and middle fingers respectively). The purposes of the task were to ensure that the subject paid attention to which scene was being shown and to minimize eye movements. In addition, we used eye-tracking to make sure subjects were fixating. Subjects were familiarised with the full non-occluded scenes in a short practice run prior to going into the scanner. This was done to increase subjects’ contextual associations and thus increase meaningful feedback when viewing the scenes with reduced information in the experimental trials.

After the experimental runs, we performed a polar angle retinotopic mapping procedure to estimate the borders of the early visual areas V1-3. This consisted of a single checkerboard wedge which started in the right horizontal meridian and rotated clockwise (12 rotations per scan, wedge angle: 22.5°, scan time: 13 min 28 sec). For 10 subjects, we also performed an eccentricity mapping procedure. This consisted of an expanding ring which started at the centre and expanded towards the periphery (8 expansions per scan, ring width increased exponentially towards the periphery, scan time: 9 min 12 sec).

#### MRI acquisition

We collected MRI data using a 3T Siemens Tim Trio System with a 12-channel head coil. We measured blood oxygen level dependent (BOLD) signals with an echo-planar imaging sequence (echo time: 30 ms, repetition time: 1000 ms, field of view: 210 mm, flip angle: 62°, 18 axial slices). The spatial resolution for functional data was 3 mm^3^. Each experimental run had 648 volumes. Retinotopic mapping consisted of 808 volumes (polar angle) or 552 volumes (eccentricity). We positioned 18 slices to maximize coverage of occipital cortex. We recorded a high-resolution 3D anatomical scan (3D Magnetization Prepared Rapid Gradient Echo, 1 mm^3^ resolution, 192 volumes).

#### MRI data processing

We corrected functional data for each experimental run and retinotopic mapping runs for slice time (cubic spline interpolation) and 3D motion (Trilinear/Sinc interpolation), temporally filtered (high-pass filtered at 6 cycles with GLM-Fourier, and linearly detrended), and spatially normalized data into Talairach space with BrainVoyager QX 2.8 (Brain Innovation, Maastricht, The Netherlands; Goebel, 2012). We used the anatomical data to create an inflated cortical surface and functional data were overlaid.

#### Voxel selection and analysis

Excessive subject movement between runs is likely to affect correspondence between voxels from one run to another. This could introduce noise into our analysis as we selected our region of interest (ROI) based on the averaged functional data of all 4 runs. As described previously (Revina et al., 2018), we calculated an alignment value for each subject by measuring Pearson’s correlation in a ROI in the visual cortex between the four functional runs. Correlations were performed in a ROI covering the early visual cortex using intensity values from an anatomical representation of the first volume of the functional data of every run. High correlations would suggest a close anatomical alignment between the 4 runs. The median alignment value across subjects was 98.08% and single subject values ranged from 77.85% to 99.31%. We excluded data from further analysis if the alignment value was below 90%, which applied to three subjects. Furthermore, we excluded any subject with chance level performance in any feedforward condition in single trial analysis (significance above chance was measured using permutation analysis with 1000 trials). The feedforward conditions have bottom-up stimulation and hence there should be a difference in activity patterns. If the scenes could not be decoded in these control conditions in a subject, we excluded them from the analysis, as it suggests that the subject might not have been fixating properly, not paying enough attention, falling asleep, and so on. It would not be meaningful to assess feedback classifier performance (or lack of) in such cases. This excluded a further five subjects. Thus, the following analyses were performed on 21 subjects.

We identified the cortical representation of the occluded region using a general linear model (GLM) contrast of the *Target* region against the *Near Surround*, as described previously in Smith & Muckli (2010). The ROI was selected from activation in V1 only. To further minimize spillover activity from neighbouring stimulated areas, we selected voxels from the ROI on the basis of the difference between *Target* and *Near* S*urround* t-values being greater than 1.

##### Analyses with extended boundary around the occluded region

To further make sure our findings of scene information in the quadrant were not due to spillover activity from the feedforward surround, we performed a separate analysis with more stringent methods of voxel selection. First of all, we selected our region of interest in BrainVoyager as the contrast of the Target mapping region being higher than both the *Near Surround* and the *Inner Border* mapping conditions. In addition, we selected voxels fitting the criteria of (*Target* - *Near Surround*) > 1 and (*Target* - *Inner Border*) > 1. This helped to restrict voxels to the more peripheral regions and to further minimize any voxels at the inner borders of the quadrant. Analysis showed the same pattern of results and significant decoding between the two scenes in all conditions except Small Bubbles Feedback and Full Feedback (average block analysis, Experiment 1 only).

Moreover, we performed another analysis using population receptive field (pRF, Dumoulin and Wandell, 2008) mapping for the subjects which had both the polar angle and eccentricity retinotopic mapping available (Expt 2: n = 4, Expt 3: n = 2, Expt 4: n = 4). Again, this was done to restrict our voxel selection to the quadrant. We only included voxels that were both within the occluded region as defined by pRF and only within our original *Target > Near Surround* ROI as defined in BrainVoyager.

### Multivariate Pattern Classification Analysis

The voxels matching all the above-mentioned criteria for each analysis were entered into the linear classifier (Support Vector Machine [SVM], using the LIBSVM toolbox in MATLAB, Chang and Lin, 2001). For classification analyses, we trained the classifier to decode between the 2 scenes in each condition. For cross-classification analyses we trained the classifier to decode between the two scenes on one condition and tested on the other. The classifier used single trial activity patterns (beta values) for training, and was then tested on either “single trial” (ST; 8 trials × 4 sequences = 32 separate trials) or “average block” (AB) activity patterns for each of the 8 trial types (average of the 4 repetitions). In other words, for the average block analysis, the training was the same (single trials of three runs, 32 trials in each run) but the testing was done on the average per stimulus condition of the fourth run. For both types of analyses, we trained the classifier on 3 of the runs and tested on the remaining run (i.e. one-run-out cross-validation).

In order to get a robust average and to test how well the classifier would perform when the labels were randomly assigned (described in more detail in Revina et al. 2018), we used bootstrapping and permutation analysis. We bootstrapped the classifier performances 10000 times for individual subjects (there were four performances for each condition for each subject due to the one-run-out method on the four runs), to estimate the single subject mean. We then bootstrapped these mean values from individual subjects 10000 times to estimate 95% confidence intervals (CIs) on the group mean. We counted classifier performances as significantly above chance (50%) if the 95% CIs did not contain chance-level performance. We used a permutation test (1000 samples) to compute differences between mean group classifier performances (reported *p* values not corrected for multiple comparisons), by shuffling the observed values across the conditions, and calculating the absolute differences between the conditions. If the observed difference was in the top 5% of the differences distribution, we considered our conditions to be significantly different from each other.

## Results

Our hypothesis is that the surround stimulation drives higher visual areas with larger receptive fields to send a contextual feedback signal to voxels in V1 responding to the occluded quadrant. We can therefore modify the surround stimulation to learn more about the nature of contextual feedback.

### Increased stimulation of the surround receptive field enhances contextual feedback

We have shown previously that scene features eliciting contextual feedback to non-stimulated V1 are not only those features located nearest to the occluded region of the image (Smith and Muckli 2010). That is, voxels contributing information to scene classification are not only found near the border of the occluder (Morgan et al., 2019). This finding suggests that scene classification in non-stimulated voxels is not only related to short-range lateral connections. Expanding on these findings, we assessed the amount of surrounding scene information required to elicit scene-relevant information in non-stimulated V1. We parametrically modulated the availability of surround information and trained the SVM classifier to decode between the two scenes using voxel patterns responding to the lower right quadrant when it was either occluded (feedback and lateral, but no feedforward information) or stimulated (feedforward, feedback and lateral information). SVM classification performance was used as an estimate of the amount of available information in the activation pattern.

When the image was occluded, scene classification in non-stimulated voxels improved with increasing availability of surrounding scene information (**Figure 2A**, left). Averaging across experiments, classification was significantly above chance once the bubbles exceeded the smallest size, except for Large Bubbles Single Trial analysis (**Table 1**). Classifier performance for the Full Feedback condition was significantly higher than the Small or Medium Bubbles conditions (Small: ST: *p* < 0.001; AB: *p* = 0.015; Medium: ST only: *p* = 0.009). Increased surround information also improved classifier performance during feedforward processing of the scenes (**Figure 2A**, right), even though voxels received identical feedforward stimulation. The Fully Visible condition was significantly higher than the other feedforward conditions (Large: AB only, *p* = 0.019; Medium: ST: *p* = 0.028, AB: *p* = 0.001; Small: ST only, *p* = 0.007; ¼: ST: *p* = 0.034, AB: *p* = 0.017). Classification performance for individual experiments is shown in **Figure 2B**.

**Figure 2.**
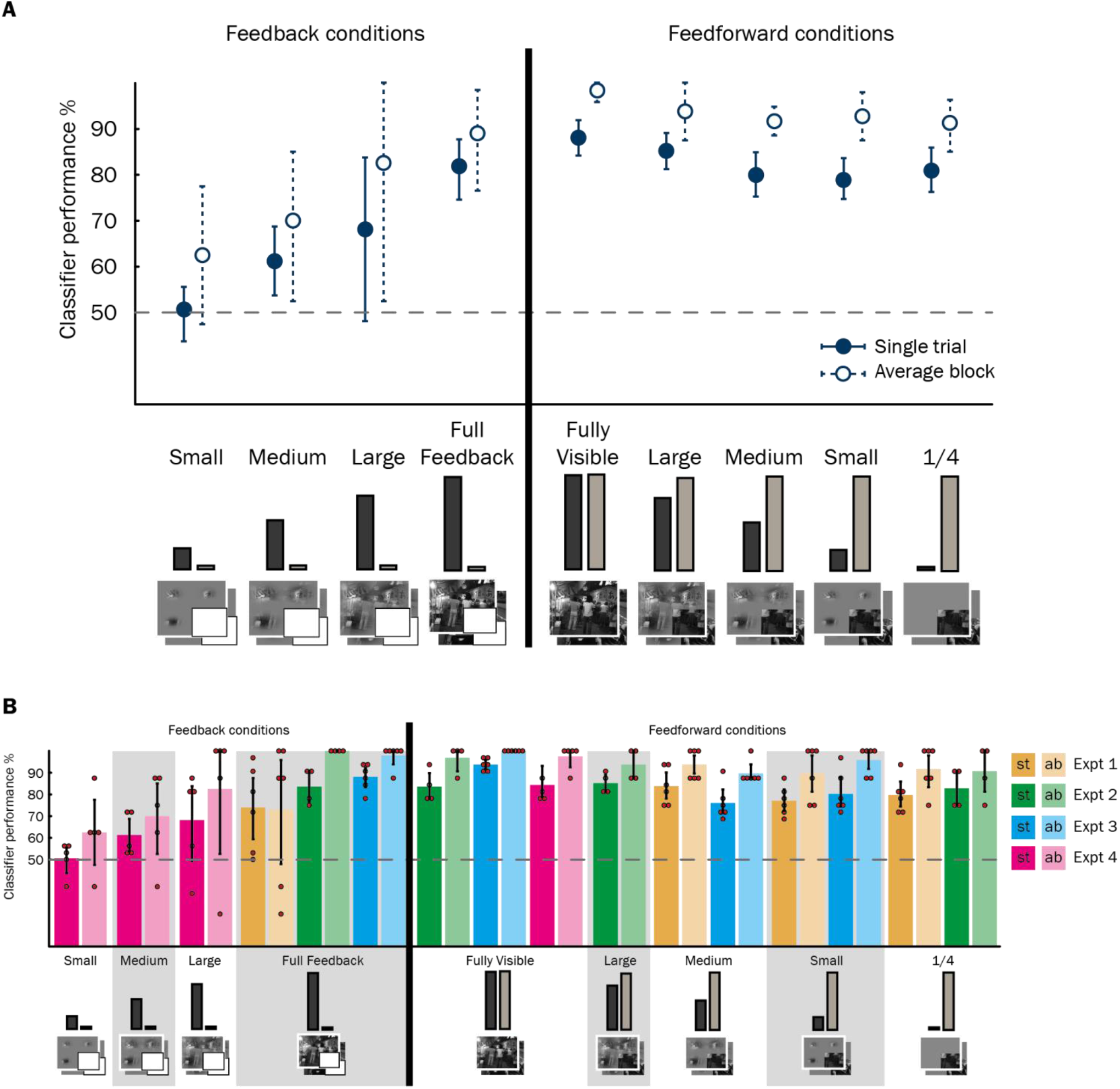
Classification performance for decoding between the two scenes in each condition, for feedback and feedforward stimuli. Chance level is 50%. Lines represent 95% confidence intervals around the bootstrapped mean (10000 bootstrap samples of individual subjects’ performances). Classifier performance is significantly above chance at α = 0.05 (not corrected for multiple comparisons) if the confidence intervals do not intersect with the chance line. A) Classifier performance for each condition, averaged over the four experiments (solid line = classifier tested on single trials; dashed line = classifier tested on blocks of conditions averaged over the same type). Small, Medium and Large Feedback conditions, n = 5; Full Feedback, n = 16; Fully Visible, n = 15; Large Feedforward, n = 4, Medium and Small Feedforward, n = 12; ¼, n = 10. B) Same data as in (A) but classifier performance split by four experiments (separate colours). ST (dark hues) show performance when classifier was tested on single trials; AB (light hues) when tested on blocks of conditions averaged over the same type. Red circles represent individual subjects’ results.

**Table 1.**
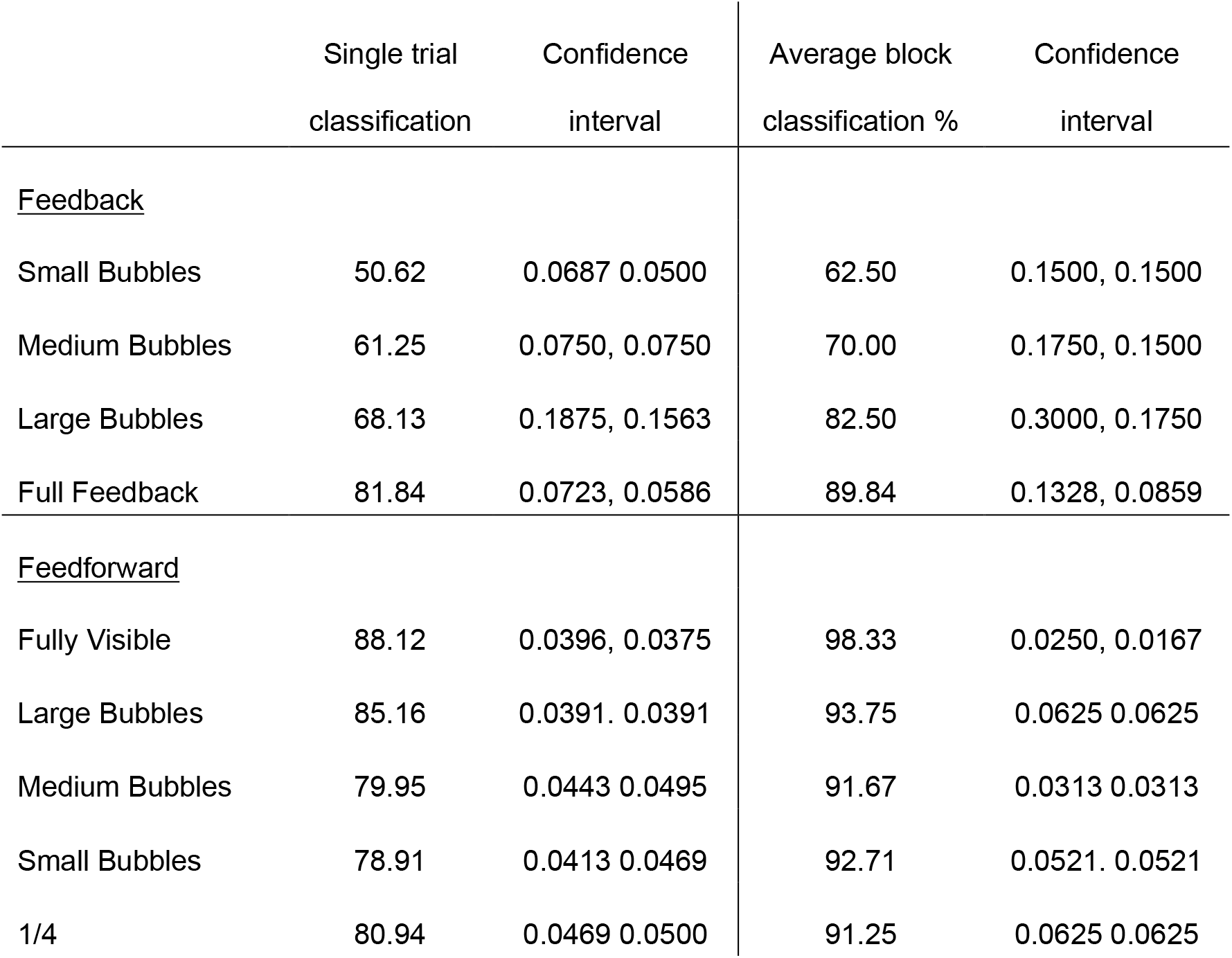
Classification performance for decoding between the two scenes in each condition, for feedback and feedforward stimuli, averaged across experiments.

|  | Single trial<br>classification | Confidence<br>interval | Average block<br>classification % | Confidence<br>interval |
| --- | --- | --- | --- | --- |
| <u>Feedback</u> |  |  |  |  |
| Small Bubbles | 50.62 | 0.0687 0.0500 | 62.50 | 0.1500, 0.1500 |
| Medium Bubbles | 61.25 | 0.0750, 0.0750 | 70.00 | 0.1750, 0.1500 |
| Large Bubbles | 68.13 | 0.1875, 0.1563 | 82.50 | 0.3000, 0.1750 |
| Full Feedback | 81.84 | 0.0723, 0.0586 | 89.84 | 0.1328, 0.0859 |
| <u>Feedforward</u> |  |  |  |  |
| Fully Visible | 88.12 | 0.0396, 0.0375 | 98.33 | 0.0250, 0.0167 |
| Large Bubbles | 85.16 | 0.0391, 0.0391 | 93.75 | 0.0625, 0.0625 |
| Medium Bubbles | 79.95 | 0.0443, 0.0495 | 91.67 | 0.0313, 0.0313 |
| Small Bubbles | 78.91 | 0.0413, 0.0469 | 92.71 | 0.0521, 0.0521 |
| 1/4 | 80.94 | 0.0469, 0.0500 | 91.25 | 0.0625, 0.0625 |

### Contextual feedback enhances feedforward processing

Classifier analyses so far reveal that increased presence of the surrounding scene enhances scene-specific information in non-stimulated V1. This finding is consistent with the hypothesis that part of V1 neuronal information patterns comprises feedback signals from areas with larger receptive fields higher up in the visual hierarchy. Interestingly, feedforward information was also enhanced with increased surround stimulation. Observing informative feedback signals with and without feedforward input motivates the question: do feedback and feedforward signals carry the same information? We used a cross-classification approach to test if the classifier can discriminate the two scenes in the feedback conditions and then use this information to discriminate scenes in the feedforward condition. Successful cross-classification would suggest similar information content in feedforward and feedback signals.

### How much does feedback contribute to visual processing?

We trained the classifier to decode between the two scenes in the Full Feedback condition (with no direct feedforward input in the quadrant) and tested on the feedforward conditions, with varying amount of feedback from the surround (**Figure 3**). Cross-classification performance decreased with decreasing scene information in the surround. The classifier could generalize from the Full Feedback condition to the Fully Visible and Large Bubbles condition (ST only; **Figure 3A** and **Table 2**). However, cross-classification for Medium, Small Bubbles, and ¼ conditions was at chance level. Averaging across experiments (**Figure 3A**), the Fully Visible condition was significantly higher than the Medium Bubbles (ST: *p* = 0.002; AB: *p* = 0.021), Small Bubbles (ST: *p* < 0.001; AB: *p* < 0.001) and the ¼ condition (ST: *p* < 0.001; AB: *p* = 0.003). These results tell us that we can train on a feedback signal (that likely has a coarser resolution of information), and test on a signal that is a combination of fine-grained feedforward signal and (coarse) surround feedback signal. This cross-classification must be due to the contextual feedback signal rather than shared information between feedforward and feedback because when the surround stimulus is reduced to nothing (i.e. with shrinking bubbles), we learn that the content of information or its scale (coarse or fine) in feedforward and feedback signals differs.

**Figure 3.**
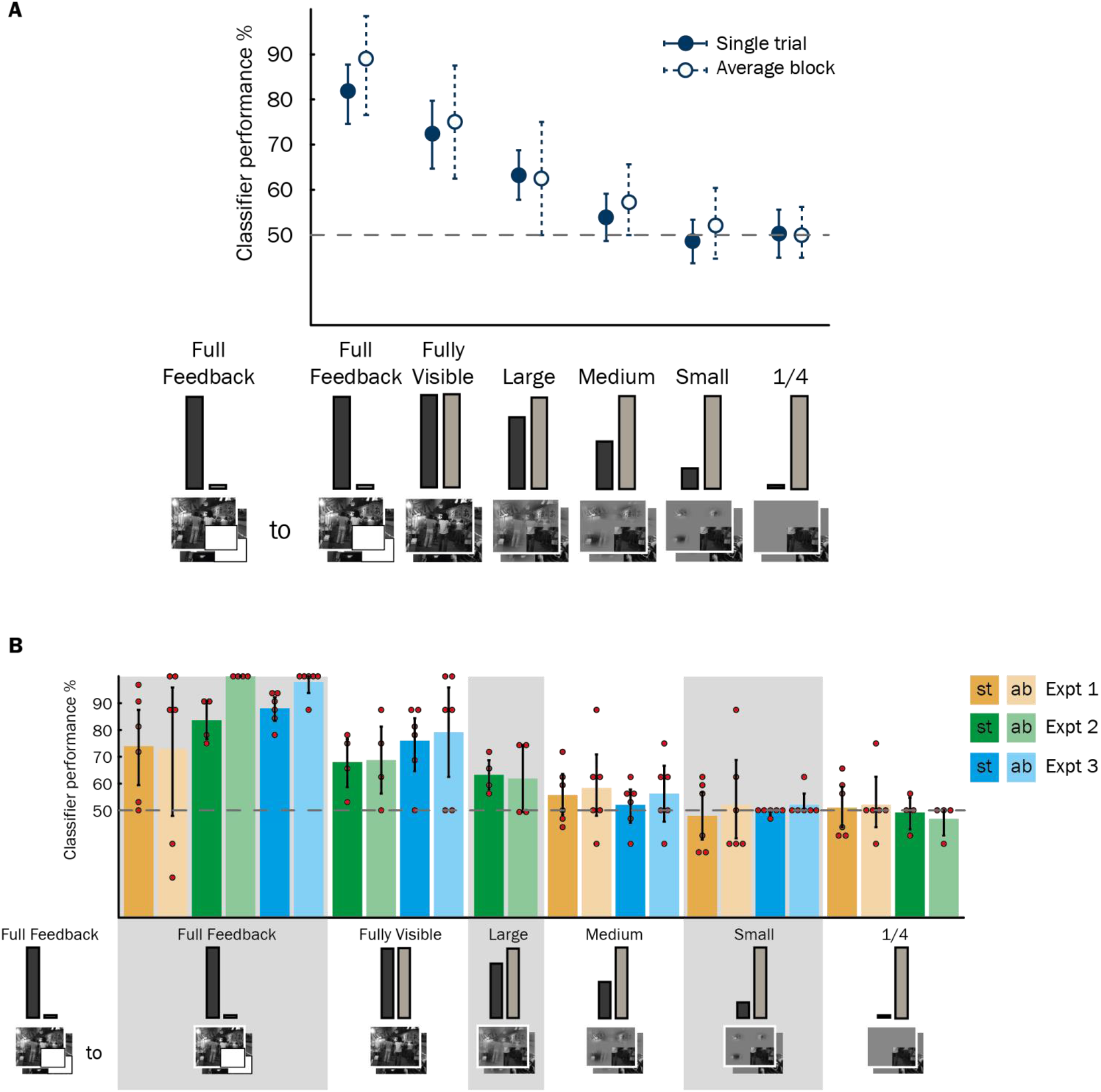
Cross-classification performance for training on the Full Feedback condition and testing on the feedforward conditions. Chance level is 50%. Lines represent 95% confidence intervals around the bootstrapped mean. Classification performance for the Full Feedback stimulus (training and testing on the same condition) is shown for comparison. Classifier performance is significantly above chance at α = 0.05 (not corrected for multiple comparisons) if the confidence intervals do not intersect with the chance line. A) Classifier performance for each condition, averaged over the four experiments (solid line = classifier tested on single trials; dashed line = classifier tested on blocks of conditions averaged over the same type). Fully Visible, n = 10; Large, n = 4; Medium and Small, n = 12; ¼, n = 10. B) Same data as in (A) but classifier performance split by four experiments (separate colours). ST (dark hues) show performance when the classifier was tested on single trials; AB (light hues) when tested on blocks of conditions averaged over the same type. The small red circles represent individual subjects’ results.

**Table 2.**
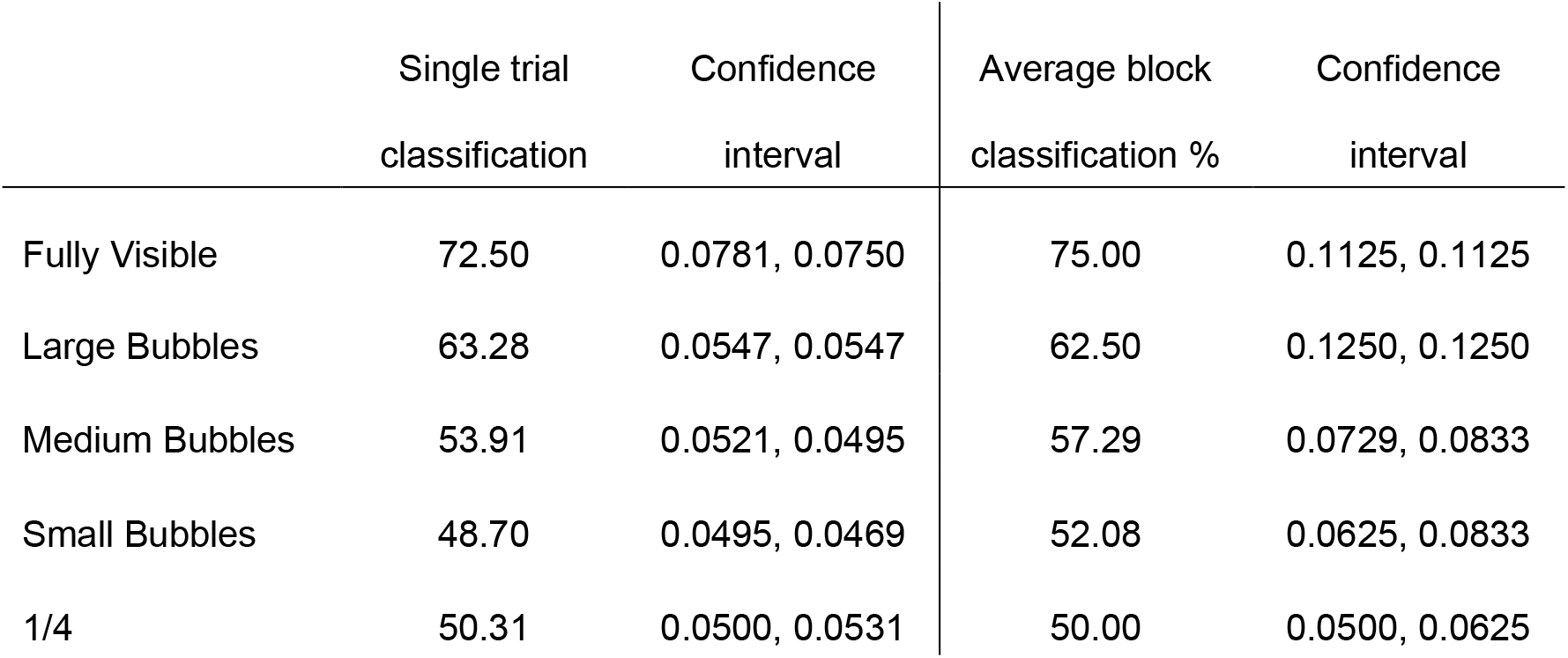
Cross-classification performance for training on the Full Feedback condition and testing on the feedforward conditions, averaged across experiments.

|  |  | Single trial | Confidence | Average block | Confidence |
| --- | --- | --- | --- | --- | --- |
|  |  | classification | interval | classification % | interval |
|  | Fully Visible | 72.50 | 0.0781, 0.0750 | 75.00 | 0.1125, 0.1125 |
|  | Large Bubbles | 63.28 | 0.0547, 0.0547 | 62.50 | 0.1250, 0.1250 |
|  | Medium Bubbles | 53.91 | 0.0521, 0.0495 | 57.29 | 0.0729, 0.0833 |
|  | Small Bubbles | 48.70 | 0.0495, 0.0469 | 52.08 | 0.0625, 0.0833 |
|  | 1/4 | 50.31 | 0.0500, 0.0531 | 50.00 | 0.0500, 0.0625 |

To further test how much surround information contributes to visual processing, we compared the Fully Visible scene with other feedforward conditions with a reduced scene surround, as well as the feedback conditions (**Figure 4**). We trained the classifier on the Fully Visible scene and tested on the other conditions. In a fully visible scene both parts of the information are available simultaneously and the classifier might rely more on the rich, fine-grained feedforward information. However, we found that Fully Visible feedforward to feedback cross-classification was only possible with large amounts of scene information surrounding the occluded region (**Table 3**). Fully Visible to Full Feedback cross-classification was above chance, while Large, Medium and Small Bubbles did not reach significance in the feedback conditions. In addition, although we could cross-classify above chance from the Fully Visible to all other feedforward conditions, cross-classification reduced with decreased scene information in the surround. Classifier performance was significantly higher for Large Bubbles compared to Small Bubbles (ST: *p* = 0.007; AB: *p* = 0.023) and ¼ (ST only: *p* = 0.028) conditions. If contextual feedback did not contribute scene-specific information to V1, we would have observed equal cross-classification across feedforward conditions, regardless of surround stimulation. This suggests that much of the information in the activity patterns of the Fully Visible scene comes from feedback from the surround.

**Figure 4.**
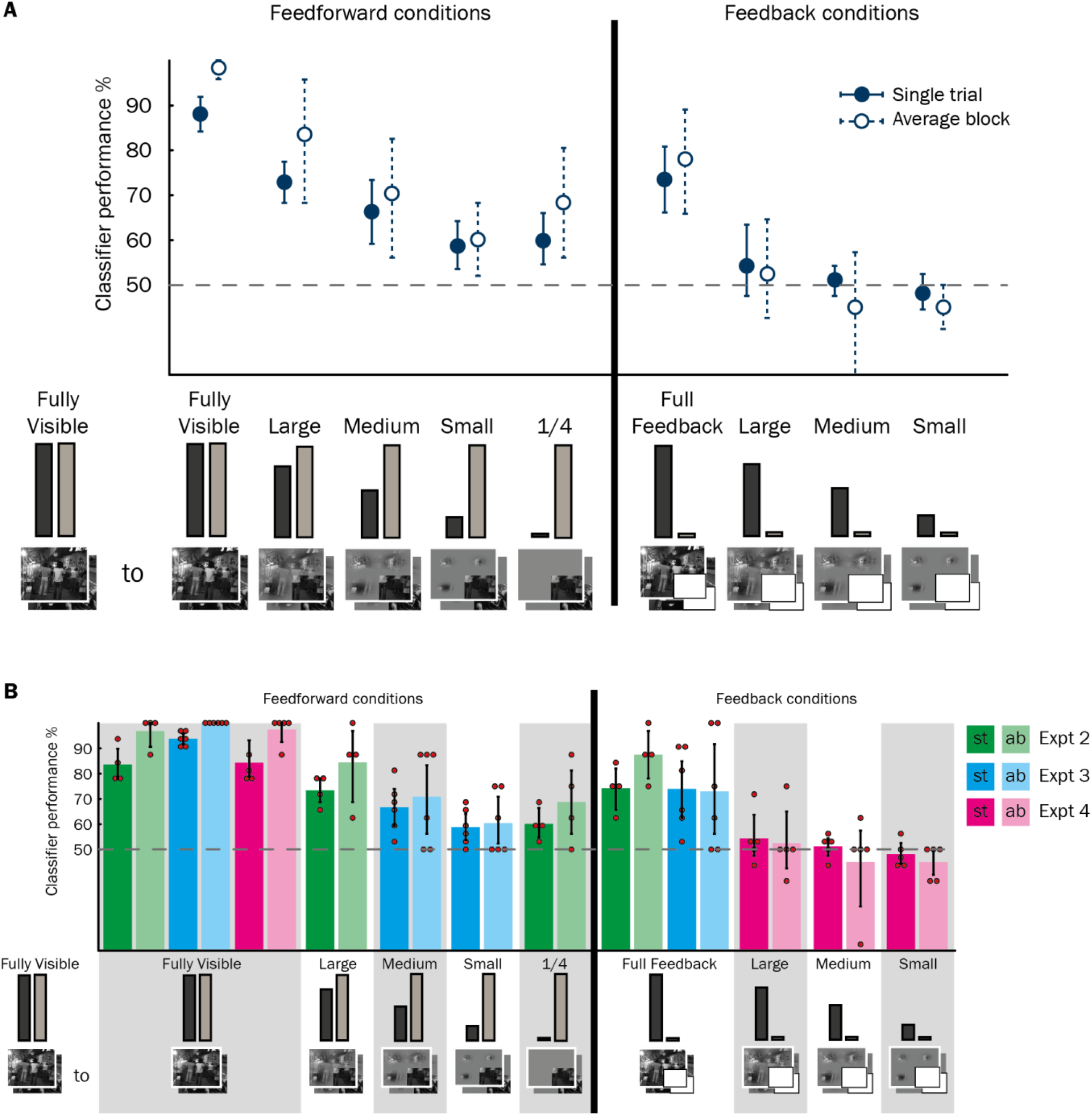
Cross-classification performance for training the classifier on the Fully Visible scene and testing on the other feedforward and feedback conditions. Chance level is 50%. Lines represent 95% confidence intervals around the bootstrapped mean. Classification performance for the Fully Visible stimulus (training and testing on the same condition) is shown for comparison. Classifier performance is significantly above chance at α = 0.05 (not corrected for multiple comparisons) if the confidence intervals do not intersect with the chance line. A) Classifier performance for each condition, averaged over the four experiments (solid line = classifier tested on single trials; dashed line = classifier tested on blocks of conditions averaged over the same type). Large Feedforward, n = 4; Medium and Small Feedforward, n = 6; ¼, n = 4; Full Feedback, n = 10; Large, Medium and Small Feedback, n = 5. B) Same data as in (A) but classifier performance split by four experiments (separate colours). ST (dark hues) show performance when the classifier was tested on single trials; AB (light hues) when tested on blocks of conditions averaged over the same type. The small red circles represent individual subjects’ results.

**Table 3.**
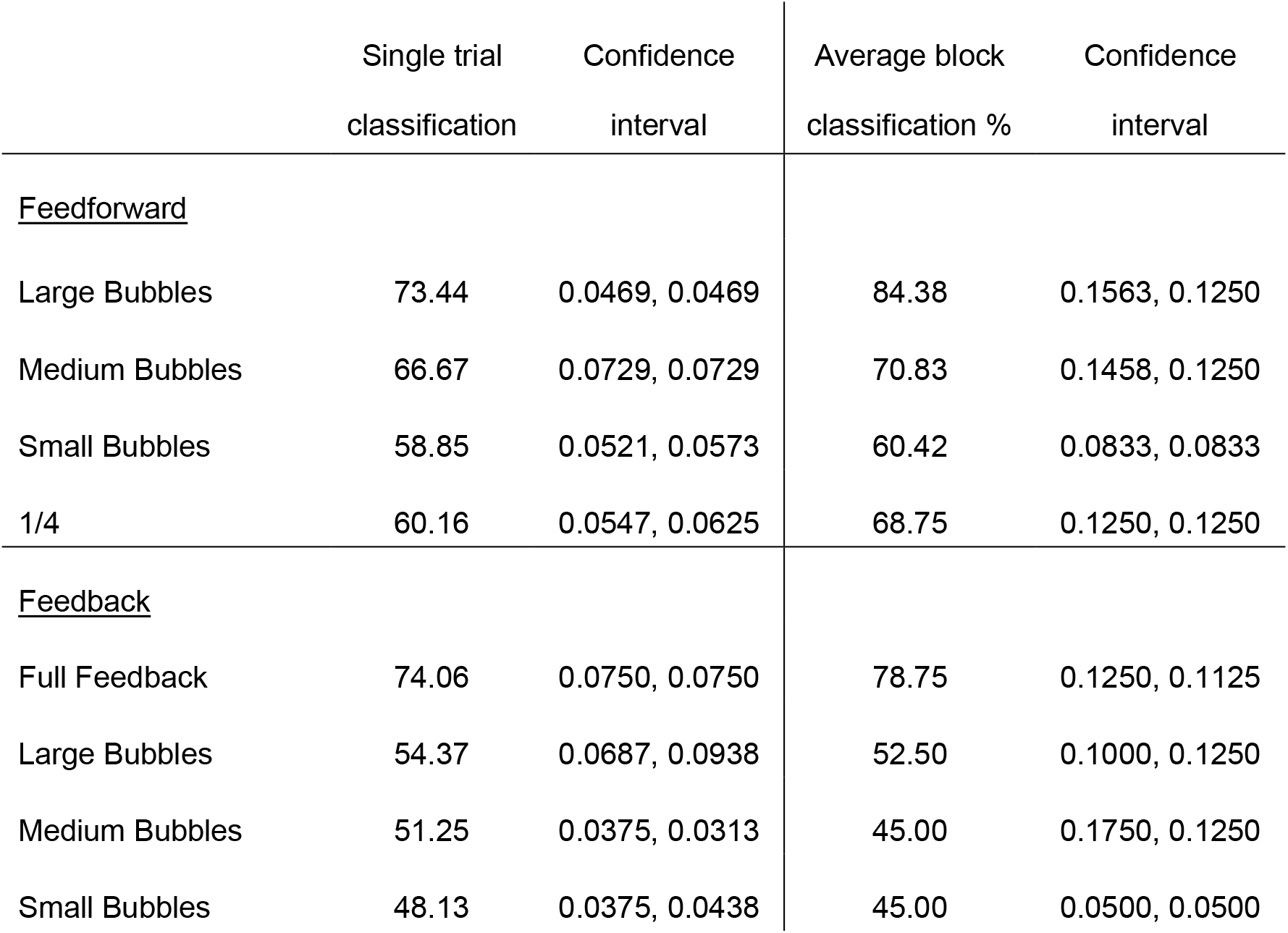
Cross-classification performance for training the classifier on the Fully Visible scene and testing on the other feedforward and feedback conditions, averaged across experiments.

Interestingly, we found that when the classifier was trained on the Fully Visible image (**Figure 4**) it cross-classified better to Full Feedback than to feedforward conditions in which the feedback was restricted by reduced contextual information (significantly above chance for Small Bubbles: ST: *p* = 0.013; AB: *p* = 0.035). This suggests that feedback in the occluded region from full stimulation in the surround is at least as informative about the scene as feedforward information in the quadrant with minimal surround stimulation. This shows that feedback is an important part of the information in V1, both when feedforward stimulation is present and when it is absent.

If surround feedback information interacts with feedforward processing, then increasing contextual surround information should reduce cross-classification from the ¼ feedforward condition to feedforward conditions with surround stimulation (**Figure 5**). Indeed, cross-classifier performance for ¼ to Small Bubbles (**Table 4**) was higher than to Large (ST only: *p* = 0.015) or the Fully Visible condition (ST: *p* = 0.021; AB: *p* = 0.006). Cross-classifier performance for ¼ to Medium Bubbles was also significantly higher than to Large (ST only: *p* = 0.039) or the Fully Visible condition (ST: *p* = 0.037; AB: *p* = 0.036).

**Figure 5.**
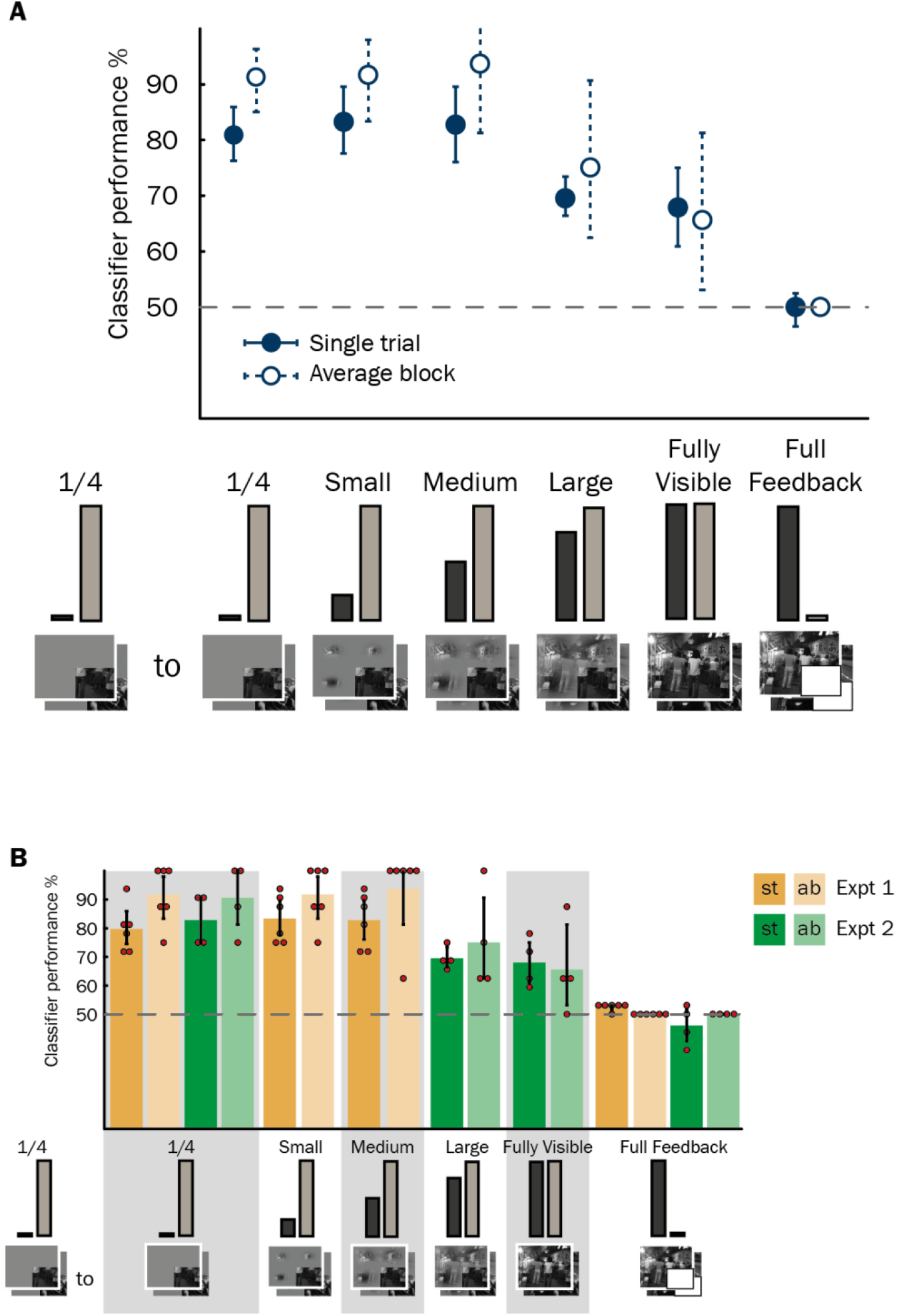
Cross-classification performance for training the classifier on the ¼ condition and testing on the other feedforward and feedback conditions. Chance level is 50%. Lines represent 95% confidence intervals around the bootstrapped mean. Classification performance for the ¼ stimulus (training and testing on the same condition) is shown for comparison. Classifier performance is significantly above chance at α = 0.05 (not corrected for multiple comparisons) if the confidence intervals do not intersect with the chance line. A) Classifier performance for each condition, averaged over the four experiments (solid line = classifier tested on single trials; dashed line = classifier tested on blocks of conditions averaged over the same type). Small and Medium, n = 6; Large and Fully Visible, n = 4; Full Feedback, n = 10. B) Same data as in (A) but classifier performance split by four experiments (separate colours). ST (dark hues) show performance when the classifier was tested on single trials; AB (light hues) when tested on blocks of conditions averaged over the same type. The small red circles represent individual subjects’ results.

**Table 4.** Cross-classification performance for training the classifier on the 1/4 and testing on the other feedforward conditions, averaged across experiments.

|  | Single trial<br>classification | Confidence<br>interval | Average block<br>classification % | Confidence<br>interval |
| --- | --- | --- | --- | --- |
| Small Bubbles | 83.33 | 0.0573, 0.0625 | 91.67 | 0.0833, 0.0625 |
| Medium Bubbles | 82.81 | 0.0677, 0.0677 | 93.75 | 0.1250, 0.0625 |
| Large Bubbles | 69.53 | 0.0313, 0.0391 | 75.00 | 0.1250, 0.1563 |
| Fully Visible | 67.97 | 0.0703, 0.0703 | 65.63 | 0.1250, 0.1563 |

### Does increased presentation of the entire image change feedback information?

Apart from varying how much surround information is visible in a stimulus, we also investigated whether knowledge of the full scene would improve feedback in the stimuli with reduced surround. In Experiment 3, we presented the Fully Visible scenes along with the Medium and Small Bubbles stimuli, unlike in Experiment 1 where we had not presented the Fully Visible scene as one of the stimuli (although all subjects were shown the full scenes in a practice run prior to the experimental session). Varying the frequency of the Fully Visible scene allowed us to investigate whether being presented with the full structure of the scene (during the experimental run) would boost meaningful feedback in stimuli with reduced surround information. We found that cross-classification from Full Feedback to Small and Medium Bubbles was at chance level for both Experiment 1 and 3 (**Figure 3B**), suggesting that reduced feedback to the feedforward quadrant in the Small and Medium Bubbles stimuli was mainly due to the decreased contextual surround information in the stimulus as opposed to a reduced implicit memory of the fully visible scene.

### Results with extended safety boundary around occluded region

We performed an additional separate analysis in order to decrease the number of voxels that are close to the boundary region and hence reduce the possibility of any feedforward stimulation “spilling over” from the surround. For the conjunction analysis using the contrast of (*Target* > *Near Surround*) & (*Target* > *Inner Border*), we found the same pattern of results and significant decoding between the two scenes in all conditions except Small Bubbles Feedback, and Full Feedback (AB analysis, Experiment 1 only).

After restricting voxels to the occluded region using pRF mapping, we saw that classifier performance decreased in some conditions, but the pattern of the results remained the same (**Figure 6**). Due to the low numbers of subjects in each experiment for whom we were able to perform pRF mapping, we did not calculate confidence intervals for some of the mean values.

**Figure 6.**
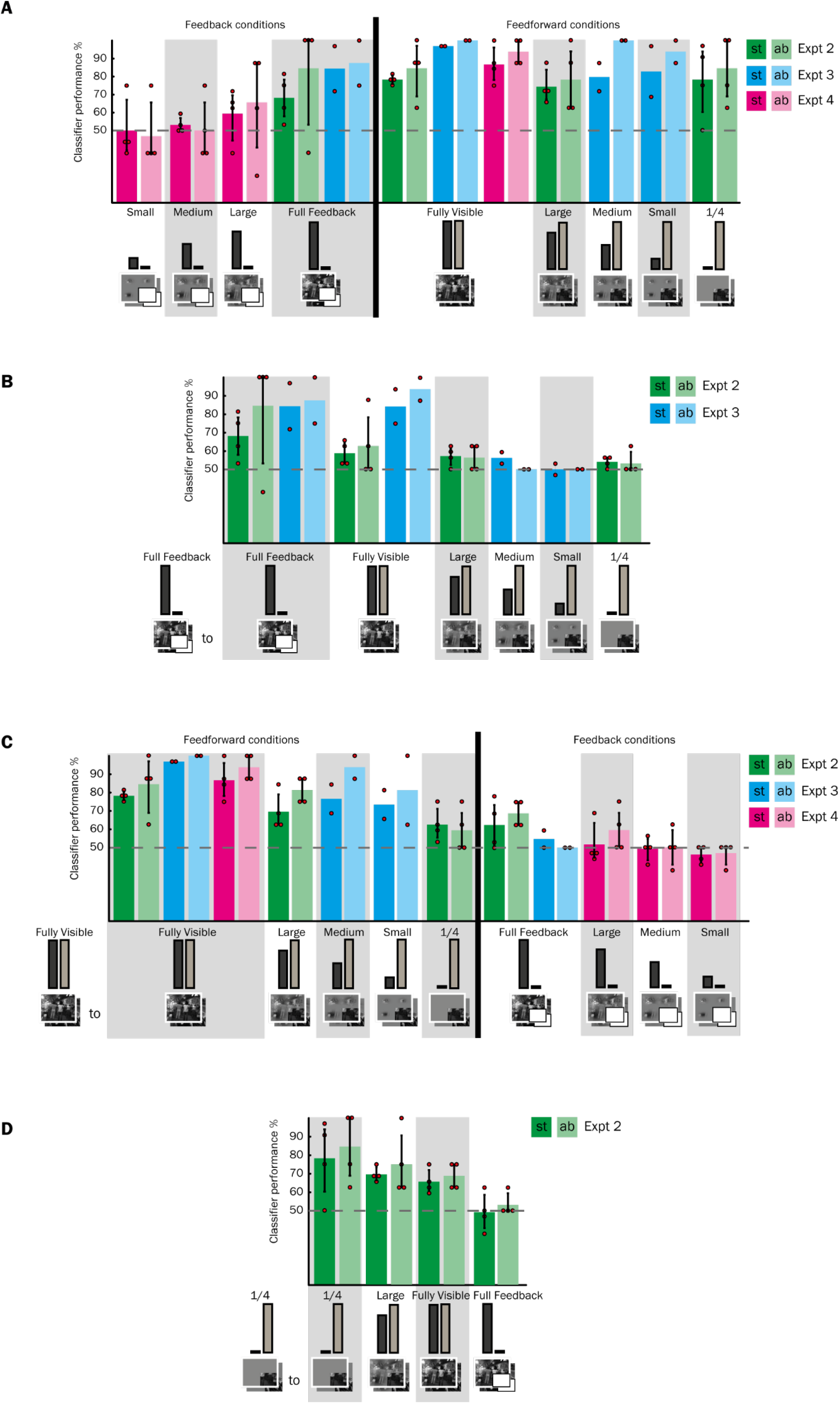
Classification and cross-classification performance after applying population receptive field (pRF) mapping to further constrain voxels to the occluded region. Classifier performance is shown for each condition for each of the three experiments (separate colours; Expt 2: n = 4; Expt 3: n = 2; Expt 4: n = 4). ST (dark hues) show performance when the classifier was tested on single trials; AB (light hues) when tested on blocks of conditions averaged over the same type. The small red circles represent individual subjects’ results. Chance level is 50%. A) Decoding two scenes in the same condition. B) Training on Full Feedback and testing on feedforward conditions. C) Training on the Fully Visible scene and testing on other feedforward and feedback conditions. D) Training on the ¼ condition and testing on other feedforward and feedback conditions.

## Discussion

We studied the influence of the scene surround on populations of neurons using fMRI, complementing what we know from electrophysiology studies investigating classical and non-classical neuronal receptive fields. We established that the availability of contextual information affects cortical feedback to a non-stimulated region in V1. Specifically, the extent of contextual modulation in non-stimulated V1 depends on the amount of scene information surrounding the occluded image quadrant. Furthermore, information in the non-stimulated region does not represent a direct filling-in of missing feedforward input, and that contextual feedback enhances information in V1 even when rich feedforward information is available.

V1 neurons integrate signals over a large area beyond the classical receptive field (Angelucci et al., 2002; Angelucci and Bressloff, 2006). Lateral connections modulate the response in the central receptive field over short distances. However, feedback from higher areas accounts for the full extent of the surround modulation effects (Angelucci and Bressloff, 2006). There was no meaningful feedforward stimulation in our occluded region of V1, and yet we could decode two scenes using information patterns corresponding to this non-stimulated region. This differential information must originate from contextual information in the scene surround. Classical receptive fields are smaller than the surround, hence neurons in the occluded area in V1 most likely receive information about the rest of the scene via cortical feedback from higher areas. Since we are measuring a population of neurons using fMRI, as opposed to single cells, it is hard to estimate how widespread the effect of the surround receptive field is. V1 receives feedback from many cortical areas, which have increasing receptive field sizes moving to higher and more abstract processing areas (Dumoulin and Wandell, 2008). Therefore, we expect that influence from the surround might be restricted to regions close to the occluded region for feedback coming from V2, for example, but transmit information from a larger area of the surrounding scene for feedback originating from higher visual areas.

We found that larger bubbles in the surround lead to more informative feedback in the occluded region. This may be because we are revealing more of the overall scene structure as we increase the bubble size. Tang and colleagues (2014) demonstrated top-down effects in image completion by presenting partially revealed images using bubbles. The number of bubbles was constant, but their location was changed. This suggests that revealing a certain amount of the global image structure, regardless of the specific parts, can be enough for top-down completion to take effect. Alternatively, our result could be explained by larger bubbles providing more information close to the lower right quadrant, compared to small bubbles, because our bubbles were centred in each quadrant. However, Williams and colleagues (2008) have demonstrated that feedback can come from distant retinotopic regions, by showing that the fovea receives feedback about objects in the periphery. Since we did not specifically measure effects of bubble location, it remains to be seen how varying proximity of surrounding information affects feedback information in non-stimulated V1. It also remains to be seen how contextual feedback depends on the presence of task-specific diagnostic features that could be revealed on different trials using bubbles (Gosselin and Schyns, 2001). Subsets of stimulus features drive information states of functionally-relevant higher brain regions (e.g. face areas) during the feedforward sweep of visual processing (e.g. Schyns et al., 2007) and these representations could modulate the content of cortical feedback signals. For the purpose of this study we kept these parameters constant.

### Interaction of feedback and feedforward signals

We found that stimulating the surround increased information in both the occluded region and when it contains feedforward information. Similarity between identical feedforward quadrants was reduced if the amount of information in the surround was increased. If feedback signals from the surround did not combine with feedforward information or only weakly modulated it, we would have seen similar activity patterns relating to the feedforward quadrant regardless of the surround. The feedforward signal has been traditionally considered the dominant signal, since it drives receptive fields, while feedback has been thought of mostly modulatory and not necessarily able to trigger spikes (Bullier, 2006; Bastos et al., 2012), but see Mignard and Malpeli, (1991). By using fMRI which is also sensitive to non-spiking activity (Logothetis, 2008; Muckli, 2010) we established that this modulation from feedback may be just as important as the spiking produced by stimuli in a bottom-up manner. fMRI is sensitive to postsynaptic inputs including the arrival of feedback onto the apical dendrites. Feedback can be combined with feedforward inputs arriving to the basal dendrites, meaning that individual neurons integrate internally-generated feedback signals with sensory-derived feedforward signals (Larkum, 2013), a process which might be a cornerstone of conscious perception (Phillips et al., 2016). Though this neuronal mechanism remains to be observed in the visual cortex, many studies support the notion that feedback to V1 is a crucial part of visual perception. For example, reducing feedback from higher areas such as V2, MT or hMT reduces the neuronal response the lower areas to visual stimulation in the centre RF, (Sandell and Schiller, 1982; Hupé et al., 1998, 2001; Schmidt et al., 2011) and in humans affects prediction in an apparent motion paradigm (Vetter et al., 2015).

### Information content of feedback

Predictive coding theories (Rao and Ballard, 1999; Friston, 2010; Clark, 2013) hypothesise that the occluded part of our scenes should be represented in non-stimulated cortex, based on the expected scene structure behind the occluder. Several authors have demonstrated that an expected or predicted stimulus evokes activity in V1 which is similar to activity elicited by actual bottom-up stimulation (e.g. Ban et al., 2013; Gavornik and Bear, 2014; Kok et al., 2014). Therefore, at first glance, it is surprising that we do not find similarity between the occluded region and the missing feedforward quadrant. This suggests information in feedback signals does not represent a direct filling-in of the missing feedforward input. However, a lack of a direct filling-in is not so counter-intuitive since participants do not report seeing the missing portion of the scene in occluded trials (i.e. they do not have a hallucination). Hence feedback and feedforward information may be coded in different formats, even though both carry information about the scene. For example, it may be that information is coarser in terms of its content because of the larger visual fields in higher visual areas or less precise retinotopically (e.g. de-Wit et al., 2012). Alternatively, feedback may provide a more abstract version of the scene. In a previous study, we have shown that feedback information is comparable to a line drawing completing the missing quadrant (Morgan et al., 2019). Finally, a difference in neural patterns could be observed because feedback and feedforward signals project to different cortical layers (Rockland and Virga, 1989; Muckli et al., 2015). Muckli and colleagues (2015) showed using high-resolution fMRI that during normal visual stimulation, feedforward information peaks in mid-layers of V1, while contextual feedback information peaks in the superficial layers. Recent data from neural network modelling also suggests that recurrent processing is not completing or filling-in the information to make it identical to the feedforward response, but rather it may function by suppressing occluders and enhancing responses to the hidden target (Spoerer et al., 2017). Recurrent networks also outperform feedforward models in identifying the occluded target stimulus, suggesting that feedback enhances feedforward processing.

If feedback signals are carrying expectations and predictions based on prior knowledge we might find that improved knowledge of the full scene structure would be important for meaningful feedback in the occluded region. However, it seems that knowledge about the particular scene being viewed is not necessary. Smith and Muckli (2010) previously found that contextual feedback in the occluded region is present even if participants never see the fully visible scene and were not familiarised with it. We also found that increased exposure to the full scene did not improve feedback in the conditions with reduced surround. Therefore, it appears that the contextual feedback we observed arises from the scene structure available in each trial, or knowledge of natural scene properties in general, but familiarity with the specific scene is not required for informative feedback signals. This could be because natural scenes have predictable scene statistics and much of the information they contain is redundant (e.g. Attneave, 1954; Barlow, 1961; Torralba and Oliva, 2003).

### Conclusion

We demonstrated that cortical feedback information forms a part of early visual cortex activity during visual stimulation. Using a brain imaging technique we have corroborated evidence from animal electrophysiology showing that stimulation in the far-surround receptive field modulates responses in the classical visual receptive field. We show that increased information in the scene surround results in increased scene information in both stimulated and non-stimulated visual field regions. We conclude that cortical feedback carries abstract internal models of natural scenes which are combined with more spatially-specific, detailed representations in primary visual cortex, and that the merging of high-level content of cortical feedback with feedforward signals should constrain our understanding of cortical function during perception.

## Acknowledgements

This work was supported by the European Research Council grant (ERC StG 2012_311751-‘Brain reading of contextual feedback and predictions’ awarded to LM) and BBSRC DTP Studentship (YR). This project has received funding from the European Union’s Horizon 2020 Framework Programme for Research and Innovation under the Specific Grant Agreement No. 720270, 785907, and 945539 (Human Brain Project SGA1-3). We thank Frances Crabbe for assistance with data collection.

